# Urobiome Analysis in Interstitial Cystitis/Bladder Pain Syndrome Reveals Nuanced Differences Associated with Localized Pain

**DOI:** 10.64898/2026.01.02.697114

**Authors:** Seth Reasoner, Jamisha Francis, Mollie Gidney, Afia Amponsah, Brendan Frainey, Andrew Schrepf, A. Grace Kelly, Anna Ryden, Leslie Crofford, Roger Dmochowski, Maria Hadjifrangiskou, Lindsey McKernan

## Abstract

**Purpose:** Interstitial cystitis/bladder pain syndrome (IC/BPS) is a prevalent chronic pain syndrome associated with functional urinary disorders. IC/BPS symptoms can be localized to the pelvic-region or have co-occurring widespread pain. Importantly, response to treatment depends on pain localization phenotype. The etiology of IC/BPS remains elusive, and whether bacteria contribute to IC/BPS pathophysiology remains uncertain.

**Materials and Methods:** We used urine samples collected from a longitudinal randomized controlled trial of individuals with IC/BPS to study the association of the urobiome and IC/BPS symptoms over time. Individuals provided urine samples at baseline, post-treatment, and at five months. We performed a secondary analysis on urine samples applying 16S rRNA sequencing and assigned bacterial taxonomy to amplicon sequence variants (ASVs) to characterize the urobiome. We then compared urobiome bacterial diversity, stability, and its association with IC/BPS symptoms over time. We also assessed the relationship between pain localization and the urobiome.

**Results:** As validation of this dataset, we noted a strong influence of menopausal status and recent urinary tract infection on the composition of the urobiome. We did not detect widespread differences in the urobiome that correlated with subjects’ pain localization or severity. Instead, we observed specific bacterial sequences that were altered in abundance in relation to symptomatology, such as reduced abundance of a *Dialister* ASV in persons with localized pelvic pain.

**Conclusions:** Together, this dataset advances our understanding of the urobiome in interstitial cystitis/bladder pain syndrome and sets the stage for future studies on the urobiome and interstitial cystitis/bladder pain syndrome symptoms.

## 1. INTRODUCTION

Interstitial cystitis/bladder pain syndrome (IC/BPS) is a chronic pain syndrome characterized by debilitating bladder pain. IC/BPS affects an estimated 4-12 million patients in the United States [1]. The characteristic bladder pain of IC/BPS often increases with bladder filling. Other symptoms of IC/BPS include urinary frequency, urgency, painful sexual intercourse, and generalized pelvic pain [2]. Symptoms can persistent and also include at times unpredictable and significant exacerbations or “flares.” The severity of symptoms in IC/BPS is associated with detrimental effects to patients’ quality of life. Frequent urination and painful symptoms negatively impact social functioning, the ability to work, and romantic relationships. The pathophysiologic cause of IC/BPS remains uncertain.

Given the proposed inflammatory nature of IC/BPS, a microbial etiology for IC/BPS has long been sought [3–5]. In fact, many of the earliest urinary microbiome studies investigated IC/BPS to ascertain whether there was a microbial contribution to IC/BPS pathophysiology [6]. Presently, it is widely accepted that IC is a non-infectious condition [7, 8]. Nonetheless, the unclear etiology of IC/BPS makes the resident microbiota of the genitourinary tract—the urobiome—a worthwhile research avenue in IC/BPS. Similarly, there has been speculation about whether urinary tract infections (UTIs) may be associated with or trigger IC/BPS onset [9, 10]. Many of the symptoms overlap between UTIs and IC/BPS, such as dysuria and urinary urgency. The temporal relationship between UTIs and IC/BPS symptom onset and/or exacerbations remains uncertain.

Several studies have investigated the resident microbiota of the bladder in IC/BPS [5, 11–15]. Another study used bladder biopsy samples to identify tissue-resident bacteria in IC/BPS [14]. Comparison of the urobiome of IC/BPS patients to healthy controls has consistently shown reduced microbiome alpha-diversity (within sample diversity) in urine samples from IC/BPS patients [5, 11, 15–17]. Conflicting results exist regarding the abundance of *Lactobacillus* in IC/BPS relative to healthy controls [5, 15, 17, 18]. These discrepancies likely arise from differences in sequencing technologies used and incomplete resolution of sub-types (i.e. species) of *Lactobacillus* [16]. Thus far, the urobiome has not revealed any conclusive association with the pathogenesis or symptomatology of IC/BPS [17, 19].

Inconsistent findings from previous investigations may be related to both sample and patient characteristics. Recent and novel research conducted by the Multidisciplinary Approach to Chronic Pelvic Pain (MAPP I/II; [20]) network emphasizes different pain “phenotypes” in IC/BPS, including individuals with more widespread versus localized pain [21]. IC/BPS phenotypes exhibit different symptom presentations, including “widespread” phenotypes demonstrating increased sites of pain the body, fatigue, energy depletion, cognitive dysfunction, and emotional distress versus individuals with “localized” pain confined to the bladder or pelvic region only without the associated non-urologic symptoms. Systemic immunologic activity has been associated with the widespread pain phenotype in IC/BPS [22]. It is possible that inconsistent and variable reports on the urobiome are related to a patient’s phenotypic presentation [19]. It is also important to examine the microbiome over time, especially given the variability of symptom severity in IC/BPS. Given similar symptoms between IC/BPS symptom flares and UTIs, it is crucial to determine whether microbial changes are linked to symptom flares to prevent potentially unwarranted antibiotic usage.

Unlike prior urobiome studies on IC/BPS, our study includes extensive clinical data regarding IC/BPS symptomatology and pain phenotype. The vast majority of urobiome studies have been cross-sectional, collecting a single sample from a patient at a single timepoint. This project aimed to evaluate the stability of the urobiome in IC/BPS and its relationship to phenotypic characteristics over time. We leveraged urine samples collected as a part of a randomized clinical trial of psychotherapy for IC/BPS to study temporal variation in the urobiome. We sought to determine whether patient symptom localization or severity can be explained by the urobiome. The strength of our dataset is its longitudinal nature, including up to 3 urine samples from the same patients over a 6-month period which allowed us to quantify metrics of the stability of the urobiome.

## 2. METHODS

### 2.1 Sample and Metadata Collection

Urine samples were collected from a subset of patients in a clinical trial evaluating cognitive behavior therapy versus active symptom monitoring for IC/BPS (NCT #04275297; [23]). Study procedures and sample collection were approved by the Vanderbilt University Medical Center IRB (IRB #193461). Eligibility was determined using the RAND Interstitial Cystitis Epidemiology (RICE) screening tool and subject self-reported IC/BPS diagnosis [1, 23]. Subjects provided written informed consent. For this project, we analyzed a subset of study participants who underwent additional in person assessment visits providing data on IC/BPS phenotypic characteristics including evaluation of the urinary microbiome. In addition to completing patient-reported outcomes via Redcap [24], participants also completed in-person study visits pre-treatment (n=38), immediately post-treatment (n=35), and at three months post-treatment (n=30).

Clean catch urine samples were collected at each in person study visit. Subjects were instructed on periurethral sterilization prior to urine collection. Urine samples were immediately stored at -80°C, with a separate portion of urine saved at room temperature for urinalysis the day of the visit. Samples were shipped on dry ice to University of California San Diego Microbiome Core for DNA extraction and sequencing.

Clinical data and symptom-based measures were collected from subject completed patient-reported outcomes, clinical data gathered the day of in-person assessment visits, and via medical record review. Menopausal status was determined from the subjects’ reported last menstrual period on the day of their assessment visit. Recent UTI was determined from the RICE screening questionnaire which asks whether the subject had a UTI or antibiotic treatment for UTI within the past three months. Patient symptom severity and localization were determined using validated questionnaires including the Genitourinary Pain Index (GUPI) and Widespread Pain Index [25–27]. Complete deidentified sample metadata is included in **Additional file 1**.

### 2.2 DNA Extraction and 16S rRNA Sequencing

DNA extraction and 16S rRNA sequencing was performed at University of San Diego Microbiome Core. Total DNA was extracted from 0.5mL of urine using the MagMAX™ Microbiome Ultra Nucleic Acid Isolation Kit (A42357, ThermoFisher) automated on a KingFisher Flex. Eight extraction blanks were included per 96-well plate of DNA extraction. Serial dilutions of the ZymoBIOMICS™ Microbial Community Standard (D6300, Zymo) were extracted in parallel with urine samples, one set per DNA extraction plate. The V4 region of the 16S rRNA gene was amplified using the 515F and 806R standardized primers [28, 29]. PCR no-template blanks were included. Paired-end 250bp reads were sequenced using the Illumina MiSeq platform. Unfiltered read counts are depicted in **Figure S1A** and **Additional file 1**.

### 2.3 Processing of 16S rRNA Sequencing Data

Raw sequence files were processed in R (version 4.4.1) using DADA2 [30]. Taxonomy was assigned to amplicon sequence variants (ASVs) with the SILVA rRNA database (version 138.1) [31]. When possible, species level assignment was made with the assignSpecies DADA2 function. ASVs are identified by the first 4 letters of their genus taxonomic assignment and an ASV label based on the ASV’s read abundance in the whole dataset (e.g. Lact_ASV1, **Additional file 2**).

Using the methodology established by KatharoSeq [32], we calculated the proportion of ASVs from the mock community serial dilutions that had the expected taxonomic composition. Allosteric sigmoidal modeling showed that samples with greater than 864 reads had >50% the expected taxonomic composition (K_1/2_, **Figure S1B**). Thus, we set this as the threshold above which we considered the sample to be positive by 16S rRNA sequencing. Critically, none of the extraction blanks had greater than this read threshold (**Figure S1A, Additional file 1**). This read threshold corresponded to between 525 and 5,250 bacterial cells of mock microbial standard added per well (**Additional file 1**). Urine samples had a median of 42,646 reads (interquartile range 25016, 57979, **Additional file 1**) before bioinformatic decontamination.

We next used the statistical program Decontam to identify potential contaminants in urine samples [33]. Negative controls included 16 extraction blanks and 4 PCR blanks. Following the methodology of Karstens et. al, we benchmarked bioinformatic decontamination using the serial dilutions of a mock microbial community of known composition (**Figure S2**) [34]. We tested various Decontam thresholds with the goal of reducing contaminating ASVs without removing the expected taxa. We used a threshold of 0.5 using the prevalence method of Decontam. Within the entire dataset, there were 1061 ASVs prior to decontamination, and 365 ASVs remained after decontamination.

### 2.4 Statistical Analysis of 16S rRNA Sequencing Data

Following bioinformatic decontamination, ASVs and their corresponding taxonomy were plotted and visualized with the R packages phyloseq and vegan packages [35, 36].

We evaluated differences in β-diversity for subject variables including UTI history, menopausal status, urinalysis results, pain localization, and GUPI scores. β-diversity was evaluated by Bray-Curtis dissimilarity indexes which were calculated with the phyloseq package’s functions *ordinate* and *distance* at the ASV level. To evaluate β-diversity differences between subject metadata variables, beta permutational multivariate analysis of variance (PERMANOVA) were computed with the vegan function *adonis2* using 9,999 permutations. For β-diversity analyses, subjects with missing metadata for the variable being tested were omitted.

We evaluated associations between subject variables (including UTI history, menopausal status, pain localization, and GUPI scores) and specific bacterial taxa at the genus and ASV levels. Differential abundance testing of ASVs and genera was conducted with Microbiome Multivariable Association with Linear Models (MaAsLin2) [37]. ASVs present at least 5% of samples (i.e., prevalence = 0.05) were included for differential abundance testing; no minimum abundance was specified. Relative abundances were log transformed within the MaAsLin2 function, and the linear “LM” model was used. Subject ID was included as a random effect in the model due to multiple samples from the same subject. We performed secondary analyses including menopausal status and prior UTI as co-variates. P-values were adjusted with the Benjamini-Hochberg correction to account for multiple comparisons. Corrected P-values are referred to as false discovery rates (FDR). We considered a MaAsLin2 FDR<0.25 as significant. For differential abundance testing, missing data were included as “NA” in the linear models.

To classify transient *versus* persistent ASVs, we considered ASVs present in at least 5 distinct subjects. No minimum abundance was set. A total of 104 ASVs were included in this analysis (**Additional file 3**). Persistence was classified as the presence of the ASV in 2/2, 2/3, or 3/3 samples from the same patient. Transience was defined as an ASV only being present in one of a subject’s samples. Persistence frequency for each ASV was the number of subjects for whom the ASV was persistent divided by the number of subjects with at least one occurrence of the ASV. For persistence analysis, we used the Wilcoxon rank-sum test to compare groups, and a P value of <0.05 was considered significant.

## 3. RESULTS

### 3.1 Study Cohort and Urobiome Characteristics

Urine samples collected included 38 unique patients across 3 timepoints, totaling 103 urine samples (**Figure 1**). Subjects contributed an average of 2.62 samples (range 1-3 samples), with 27 subjects (71%) completing all three timepoints. Within this cohort there were 36 females and 2 males (Additional file 1).

**Figure 1.**
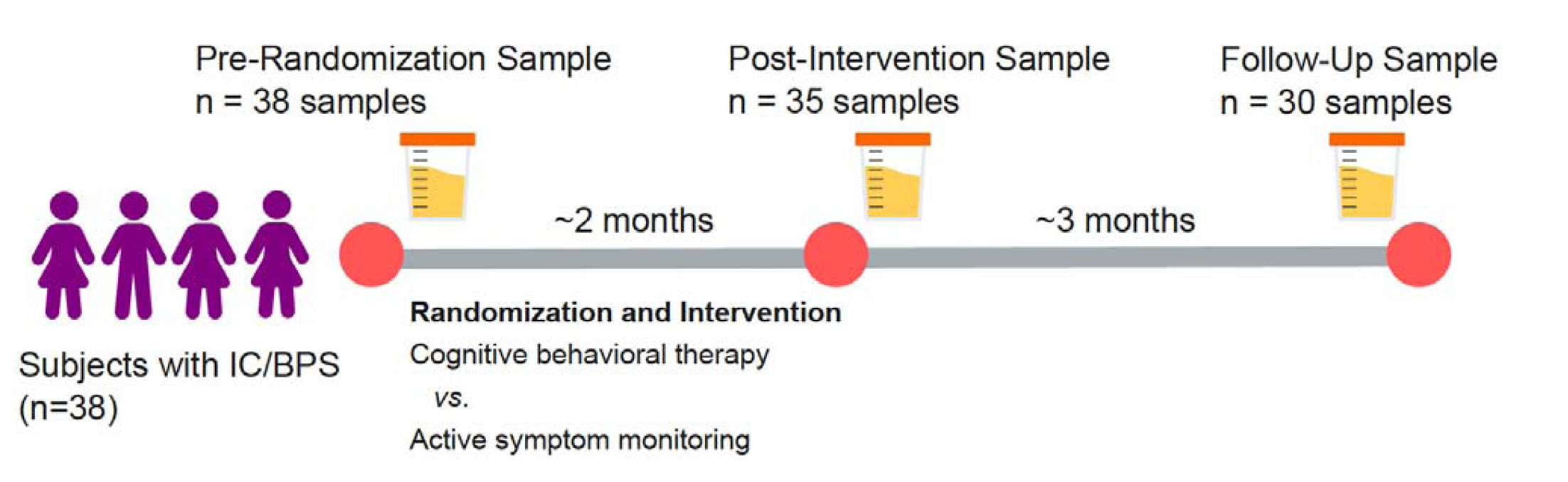
Interstitial Cystitis: Study schematic. Urine samples were collected from patients undergoing a randomized control trial for cognitive behavioral therapy (CBT) for IC/BPS. Urine samples were collected at 3 timepoints throughout the study: before randomization (time=0), after intervention (∼2 months), and at follow-up (∼5 months).

Ninety-eight percent of samples (101/103) had >864 16S rRNA reads and were classified as sequence positive. Sequencing results were bioinformatically decontaminated using Decontam (**Figure S2**). Following decontamination, the most abundant genera were *Lactobacillus* (median relative abundance, 72.4%; interquartile range [IQR], 6.4% to 94.9%), *Prevotella* (median, 1.8%; IQR, 0.05% to 11.6%), *Peptoniphilus* (median, 0.29%; IQR, 0% to 0.96%), and *Anaerococcus* (median, 0.24%; IQR, 0.02% to 1.0%). The most prevalent genera were *Lactobacillus* (94/101 samples), *Prevotella* (82/101 samples), *Anaerococcus* (77/101 samples), and *Finegoldia* (77/101 samples). Urinary pathogens such as the genera *Escherichia* and *Klebsiella* were infrequent in the dataset (present in 20 and 7 samples, respectively).

We first sought to evaluate the internal validity of our dataset. Using accompanying clinical metadata, we validated expected differences in urobiome composition and diversity. We observed significant differences in β-diversity (a metric of sample similarity or dissimilarity) between samples from pre-menopausal and post-menopausal women in the cohort (**Figure 2A**, PERMANOVA<0.001, R^2^ = 0.053). Differential abundance analysis with MaAsLin2 showed an increased abundance of a *Lactobacillus* ASV (Lact_ASV1) in pre-menopausal subjects, whereas post-menopausal subjects had an increased abundance of an *Actinotignum* ASV (Acti_ASV18), an *Enterococcus* ASV (Ente_ASV22), and an *Aerococcus* ASV (Aero_ASV122) (FDR<0.25, **Figure 2B**). Lact_ASV1 was assigned as *Lactobacillus iners*. When aggregated at the genus level, *Lactobacillus* was not differentially abundant with respect to menopause, suggesting that *L. iners* (Lact_ASV1) was reduced in menopause but other *Lactobacilli* were not uniformly reduced. This emphasizes the importance of granular taxonomic analysis. A reduction in *Lactobacillus* following menopause is well-documented for both the urinary and vaginal microbiota [38, 39]. Similarly, increased *Aerococcus* following menopause has been previously described [39]. *Enterococcus* has been reported to be present in nearly 60% of urine samples from post-menopausal women [40, 41].

**Figure 2.**
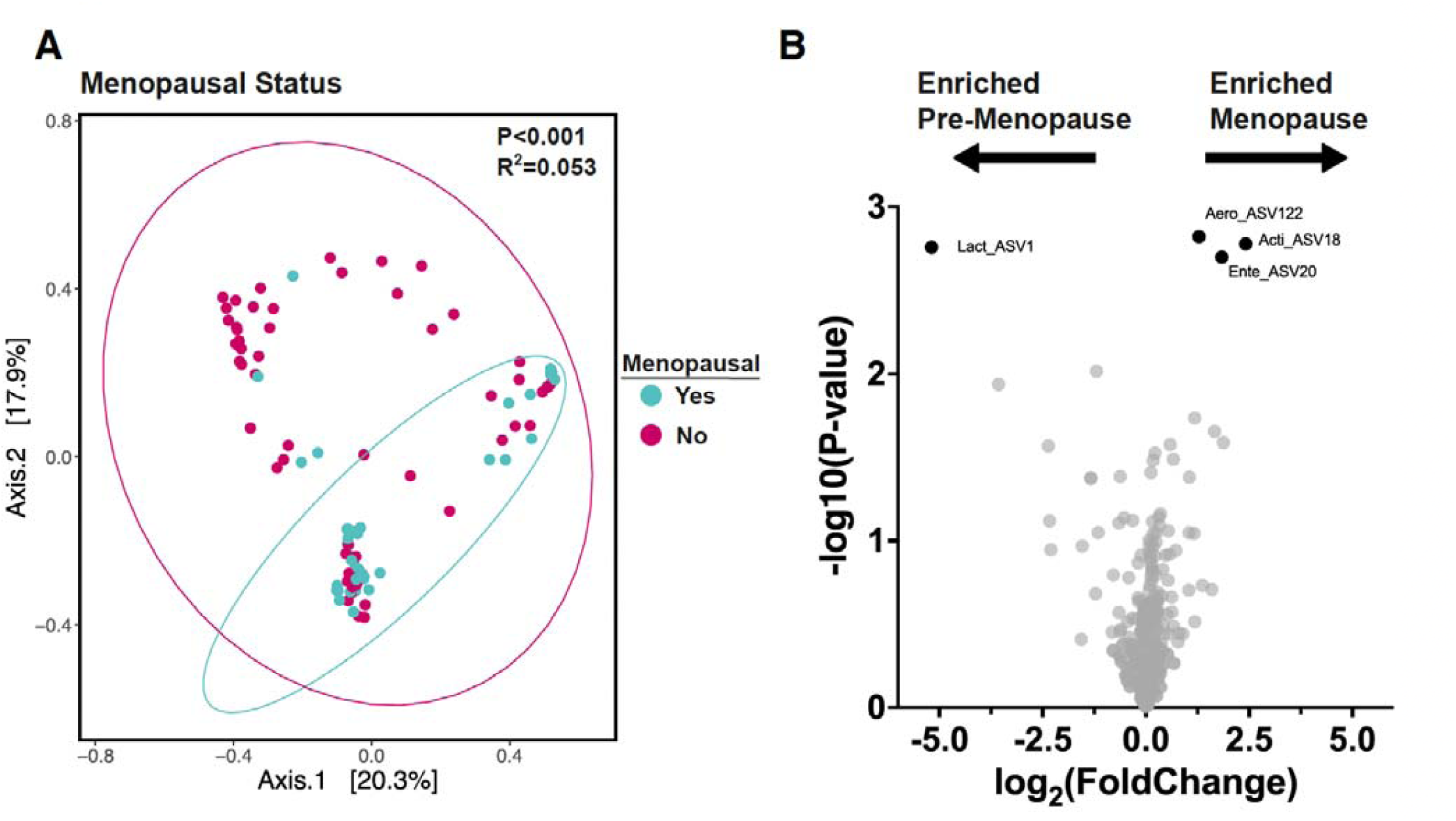
Menopause Significantly Impacts Urobiome Composition in IC/BPS. **A)** Principal component analysis of Bray-Curtis distances of samples stratified by menopausal status. P value calculated by PERMANOVA. **B)** Volcano plot of ASVs with respect to menopausal status. Differential abundance analysis conducted with MaAsLin2. Black labeled dots were significantly differentially abundant (MaAsLin2 FDR<0.25).

Urine samples from subjects who reported a urinary tract infection (UTI) within three months prior to study enrollment also had significantly different β-diversity from subjects without recent UTI (**Figure 3A**, PERMANOVA<0.001, R^2^ = 0.039). Recent UTI was associated with nominally reduced Shannon index, a measure of alpha diversity (P=0.095, **Figure 3B**). Differential abundance analysis with MaAsLin2 did not show any differentially abundant individual ASVs. At the genera level, five genera were differentially abundant with respect to recent UTI (MaAsLin2 FDR<0.25, **Figure 3C**). The genera *Peptoniphilus*, *Campylobacter*, *Lawsonella*, and *Ezakiella* were reduced in abundance in samples from subjects with recent UTI, whereas *Staphylococcus* was increased in abundance in samples from subjects with recent UTI. We also detected differences in the urobiome composition based on urinalysis results (**Figure S3**). For example, a *Veillonella* ASV was associated with the presence of nitrites on urinalysis. *Veillonella* is known to metabolize nitrates to nitrite [42]. Intrigui gly, among canonical urinary pathogens, only a *Klebsiella* ASV was associated with positive urinalysis results (**Figure S3**).

**Figure 3.**
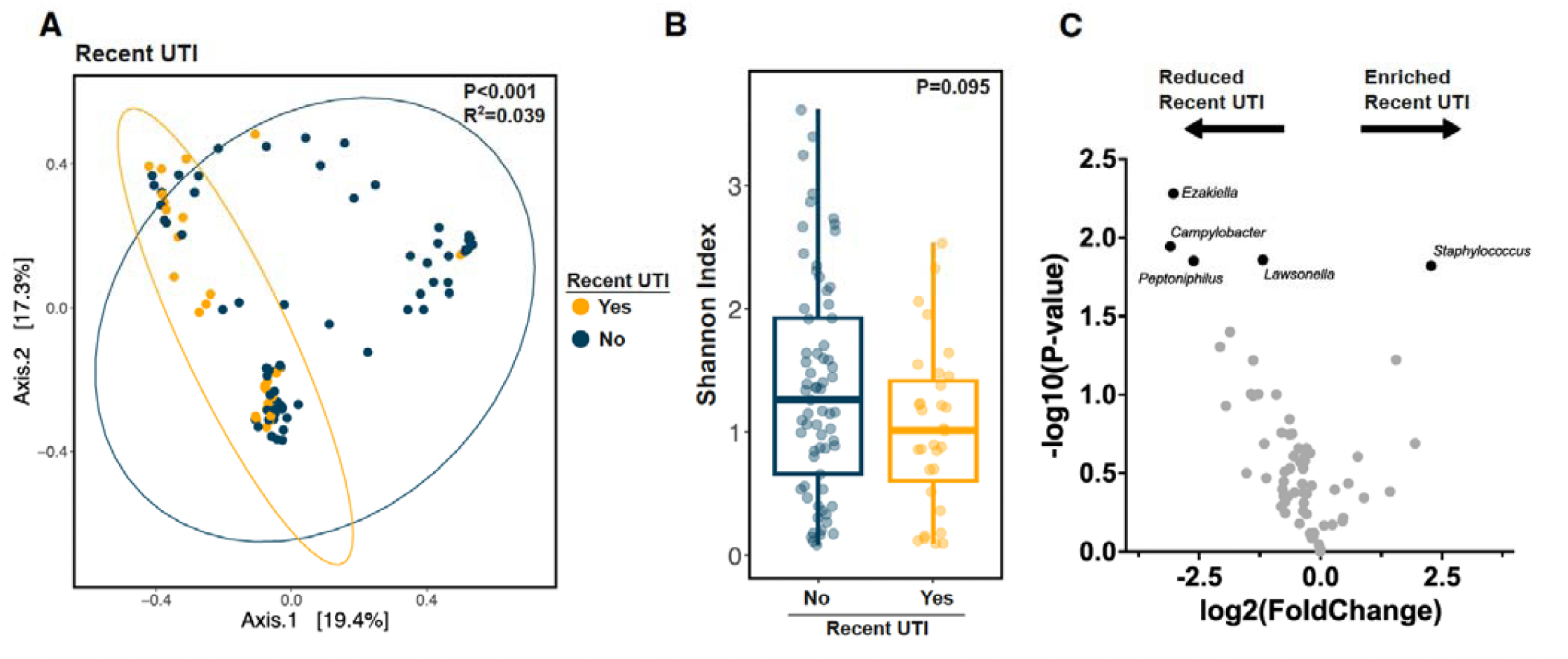
Recent UTI Impacts Urobiome Composition in IC/BPS. **A)** Principal component analysis of Bray-Curtis distances of samples stratified by recent UTI. P value calculated by PERMANOVA. **B)** Shannon alpha diversity index of samples stratified by recent UTI. P values calculated by Wilcoxon rank-sum test. **C)** Volcano plot of ASVs with respect to recent UTI. Differential abundance analysis conducted with MaAsLin2. Black labeled dots were significantly differentially abundant (MaAsLin2 FDR<0.25).

### 3.2 Association of the Urobiome with Patient Symptomatology

Per our analysis plan, we sought to evaluate the relationship between urinary microbiome and patient characteristics pain phenotype, overall symptom burden, pelvic pain severity, and urinary symptom severity. Informed by previous investigations [23, 43], we used the widespread pain index (WPI) to determine whether associations existed between patient phenotypic symptoms and the urobiome. We divided patients into phenotypic pain groups, e.g. widespread (>=3 pain sites outside the pelvic), intermediate (<3 pain sites outside the pelvic), or pelvic-only pain (no non-pelvic pain sites) based on their total WPI score [27].

We began by assessing whether symptom localization is associated with urobiome diversity or composition. Pain localization did not significantly influence overall microbiome composition (PERMANOVA=0.11, **Figure 4A**). Because some patients experienced both pelvic pain only and widespread pain at different time points, we further analyzed the subset of patients with a consistent pain phenotype across study time points (n=16 subjects). Similarly, there was no difference in overall microbiome composition between patients with consistent pain phenotypes (PERMANOVA=0.3, **Figure 4B**). As with symptom localization, overall symptom severity from the Genitourinary Pain Index (GUPI) did not correlate with overall urobiome composition (PERMANOVA=0.25, **Figure 4C**).

**Figure 4.**
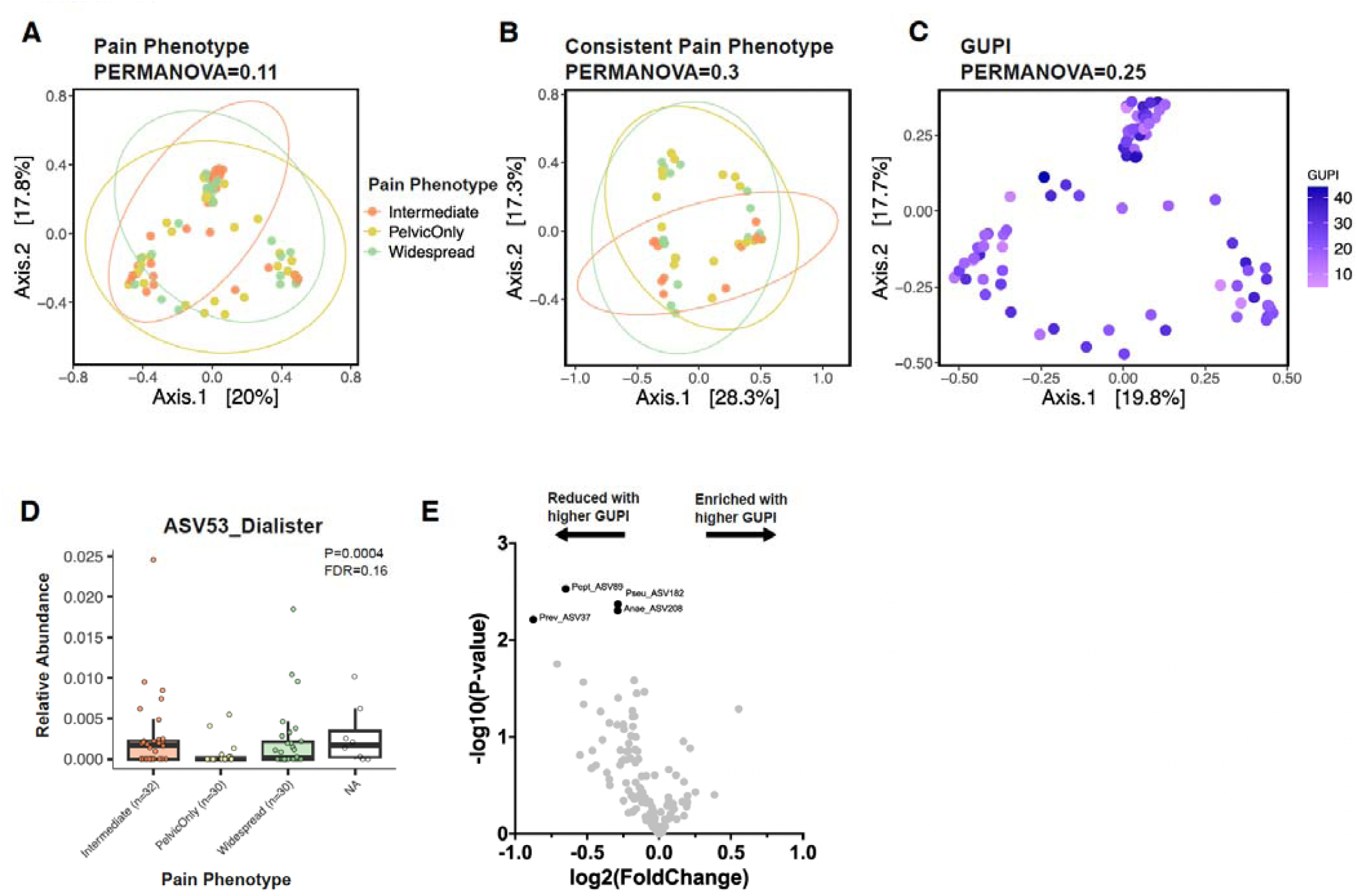
Association of Patient Symptomatology with the Urobiome. **A)** Principal component analysis of Bray-Curtis distances of samples stratified by pain localization phenotype. P value calculated by PERMANOVA. **B)** Principal component analysis of Bray-Curtis distances of samples from patients with a consistent pain localization across timepoints. Samples stratified by pain localization. P value calculated by PERMANOVA. **C)** Principal component analysis of Bray-Curtis distances of samples stratified by Genitourinary Pain Index (GUPI). P value calculated by PERMANOVA. **D)** Abundance of Dial_ASV53 with respect to pain localization. Differential abundance analysis conducted with MaAsLin2. **E)** Volcano plot of ASVs with respect to overall GUPI score. Differential abundance analysis conducted with MaAsLin2. Black labeled dots were significantly differentially abundant (MaAsLin2 FDR<0.25).

We next sought to determine whether any specific bacterial taxa were differentially abundant with respect to IC/BPS pain localization or severity of symptoms. A single ASV from the genus *Dialister* (Dial_ASV53) was reduced in abundance in patients with a pelvic-only pain phenotype (MaAsLin2 FDR=0.17, **Figure 4D**). Dial_ASV53 remained significantly less abundant in pelvic-only pain when controlling for menopausal status. Four ASVs were reduced in abundance with greater overall GUPI scores (ASV37, ASV89, ASV182, ASV208, all MaAsLin2 FDR<0.25, **Figure 4E**). These ASVs corresponded to the genera *Prevotella*, *Peptoniphilus*, *Pseudoglutamicibacter*, and *Anaerococcus,* respectively (**Figure 4E**, **Additional file 6**). We also conducted differential abundance testing using the GUPI urinary sub-score. We noted that ASVs 37, 53, 89, and 208 were among the ten most differentially abundant with respect to the GUPI urinary sub-score, though none of these ASVs met the significance threshold (**Additional file 7**, FDR>0.25). Similar differentially abundant taxa between pain phenotypes, overall GUPI score, and GUPI urinary sub-score may indicate a common pattern of bacterial taxa influencing symptomatology in IC/BPS. Importantly, none of the ASVs that were correlated with symptomatology were differentially abundant with respect to menopause status or recent UTI, suggesting that these differentially abundant ASVs are not being confounded by menopause or recent UTI.

### 3.3 Stability of the Urobiome Over Time

We evaluated the stability of the urobiome within individual patients. While 16S rRNA amplicon sequencing does not necessarily allow species or strain level taxonomic assignment, we used ASV comparison as a proxy for feature sharing between urine samples from the same subject. Between any two samples from separate patients, the average proportion of shared ASVs was 0.17 (**Figure 5A**). Between the same patient across timepoints, the proportion of shared ASVs was 0.38 (P<0.0001, **Figure 5A**). There was no difference in shared ASVs between consecutive and non-consecutive samples from the sample subject (**Figure 5B**).

**Figure 5.**
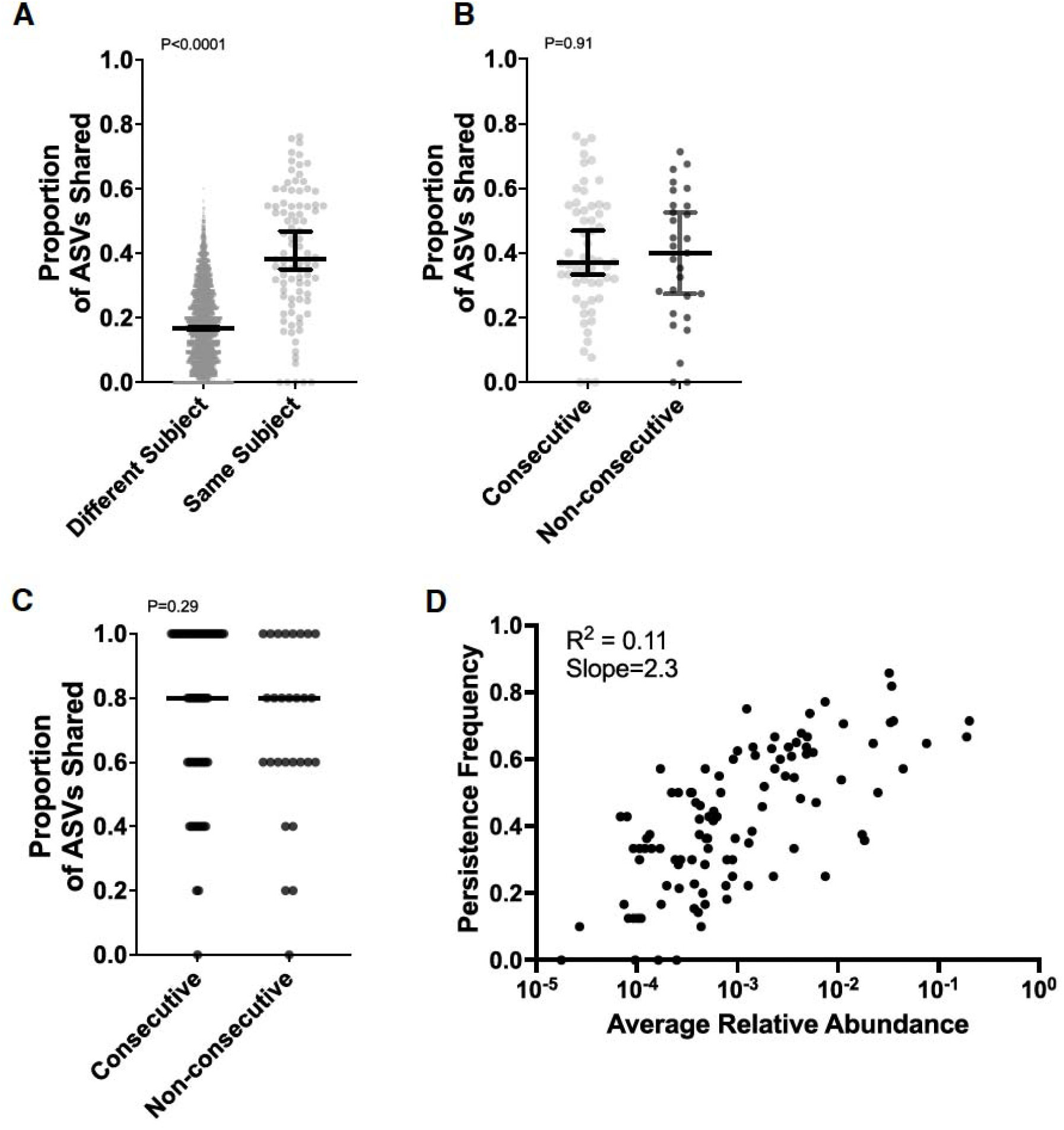
Urobiome Stability Across Timepoints and Within Subjects. **A)** Proportion of shared ASVs between samples from the same or different subjects. Error bars represent the median +/- the 95% confidence interval. P value calculated by Wilcoxon rank-sum test. **B)** Proportion of shared ASVs within the same subject at consecutive or non-consecutive timepoints. Error bars represent the median +/- the 95% confidence interval. P value calculated by Wilcoxon rank-sum test. **C)** Proportion of the top five ASVs (sorted by mean relative abundance for each subject’s samples) shared between consecutive and non-consecutive samples from the same subject. The horizontal line depicts the median. P value calculated by Wilcoxon rank-sum test. **D)** Persistence frequency vs. average ASV abundance. R^2^ value determined by simple linear regression in GraphPad Prism v. 10.2.2.

We speculated that more abundant ASVs would be more likely to be shared across timepoints. We extracted the top five ASVs from each patient’s samples by mean relative abundance and determined whether these ASVs were shared across timepoints. Indeed, among the top five ASVs from each patient’s samples, a median proportion of 0.8 were shared across timepoints (**Figure 5C**). This proportion is double that of all ASVs shared within the same patient (0.8 vs. 0.38). In fact, the most abundant ASV from each subject was nearly universally present across timepoints from the same subject (n=32/35 patients with samples from multiple timepoints).

Beyond the top five ASVs for each subject, we sought to determine which ASVs were persistent across samples from the same subject. We classified persistent ASVs as being present in >50% of samples from the same subject, whereas transient ASVs were present in a single sample from a subject. We defined the persistence frequency as the number of subjects for whom the ASV was persistent divided by the number of subjects for whom the ASV was present at least once. Across all 104 ASVs included in the persistence analysis, the mean persistence frequency was 0.42. The most persistent ASV was a *Lactobacillus* ASV (Lact_ASV9) which was persistent in 12 of the 14 subjects in whom the ASV was detected (0.86 persistence frequency). A *Staphylococcus* ASV (Stap_ASV5) was persistent in 27 of the 33 subjects in whom it was detected (0.82 persistence frequency). Comparisons of persistence frequency may allow for ecological insights to be garnered about the role of specific taxa in the urobiome. For example, all seven *Lactobacillus* ASVs had a persistence frequency >0.6, whereas ASVs assigned to the genera *Escherichia* and *Enterococcus*—known urinary pathogens—had persistence frequencies <0.4. We noted a moderate significant association between the persistence frequency and the average relative abundance of the particular ASV (R^2^= 0.11, P=0.0005, **Figure 5D**). We interpret these data as representing that the more abundant taxa are stable across timepoints whereas less abundant taxa are more likely to be transient.

## 4. DISCUSSION

Multiple factors may be contributing to IC/BPS etiology; contributing factors are thought to include urothelial dysfunction, chronic inflammation, psychosocial factors, and pelvic floor dysfunction [44]. Whether microbes contribute to IC/BPS development or pathophysiology has been a perpetual question. Our results argue against a substantial microbial contribution to pain phenotypes or symptom severity in IC/BPS. Instead, we detected specific bacterial taxa that were associated with pain localization and severity in IC/BPS. For example, an ASV corresponding to the *Dialister* genus (Dial_ASV53) was reduced in abundance among patients with a pelvic-only pain phenotype (**Figure 4D**). Intestinal abundance of *Dialister* has previously been identified to be depleted in patients with depression and negatively correlated with knee pain, suggesting a potential connection between this microbial taxa and pain [45, 46]. These taxa require validation in independent studies yet offer intriguing directions for future research regarding specific urobiome members and IC/BPS symptomatology. Together, these results suggest that large differences in urobiome composition do not determine IC/BPS symptomatology (**Figures 4A-C**), whereas specific bacterial groups may correlate with symptom localization and severity. Few urobiome studies have considered the relationship between symptoms or pain localization and the urobiome [17, 19]. Nickel et al. showed that IC/BPS subjects had increased fungal members of the urobiome during symptoms flares [47]. Similarly, a study of subjects with Hunner’s lesions showed that female subjects had no differentiating urobiome features, while men with Hunner’s lesion had increased abundance of *Negativicoccus succinivorans*, *Porphyromonas somerae*, and *Mobiluncus curtisii* and decreased abundance of *Corynebacterium renale* [48]. These prior studies are consistent with our findings that substantial urobiome changes are not correlated with symptomatology, whereas the abundance of individual microbial members may be altered in abundance.

An association between UTIs and the development of IC/BPS has been previously postulated [9]. That is, some researchers have proposed that UTIs may be a trigger for the development and/or exacerbation of IC/BPS symptoms. While our study cannot illuminate the causal relationship between UTIs and IC/BPS, we did detect a difference in urine samples from subjects with and without recent UTI (**Figure 3A**). The genera *Peptoniphilus*, *Campylobacter*, *Lawsonella*, and *Ezakiella* were increased in samples from patients with recent UTI, while *Staphylococcus* was decreased. Notably, these differentially abundant taxa are not canonical UTI pathogens. Whether these differentially abundant taxa represent direct changes from UTI or are the indirect result of antibiotic treatment cannot be determined from this dataset. These differentially abundant taxa in recent UTI samples may inform future studies regarding urobiome members and UTI pathogenesis.

We also computed metrics of urobiome dynamics. Since the vast majority of urobiome studies have been cross-sectional, little is known about the dynamics of the urobiome within individual subjects. *Lactobacillus* ASVs were nearly twice as likely to be persistent compared to *Escherichia* and *Enterococcus* ASVs. Other urobiome studies have reported that *Lactobacilli* are stable within the same subject across timepoints [49]. While broad conclusions cannot be derived from this dataset, this approach offers a framework for determining metrics of urobiome stability.

This study has notable limitations. Only one subject in the study had documented Hunner’s lesions, limiting our ability to identify potential urobiome associations with these macroscopic bladder lesions. Next, unlike prior urobiome studies of IC/BPS, our study had no healthy control population. Another limitation of this study is the lack of sampling controls collected contemporaneously with the urine samples. We implemented negative controls throughout sample processing, such as DNA extraction and PCR amplification. The orders of magnitude difference in sequencing reads between urine samples and extraction controls (**Figure S1A**) supports the authenticity of the bacterial signatures detected in urine samples. Similarly, we were able to detect differences in subject factors known to be associated with urobiome composition (i.e. menopause, recent UTI).

## 6. Conclusion

In conclusion, we used longitudinally collected urine samples to study in the intersection of the urobiome and patient symptoms in IC/BPS. Widespread differences in urobiome diversity or composition were not associated patient symptom localization or severity. We detected reduced abundance of a *Dialister* ASV in subjects with pelvic-only pain. Our future studies will attempt to independently replicate these findings and uncover the mechanisms underlying symptomatology in IC/BPS.

---

**Figure S1.**
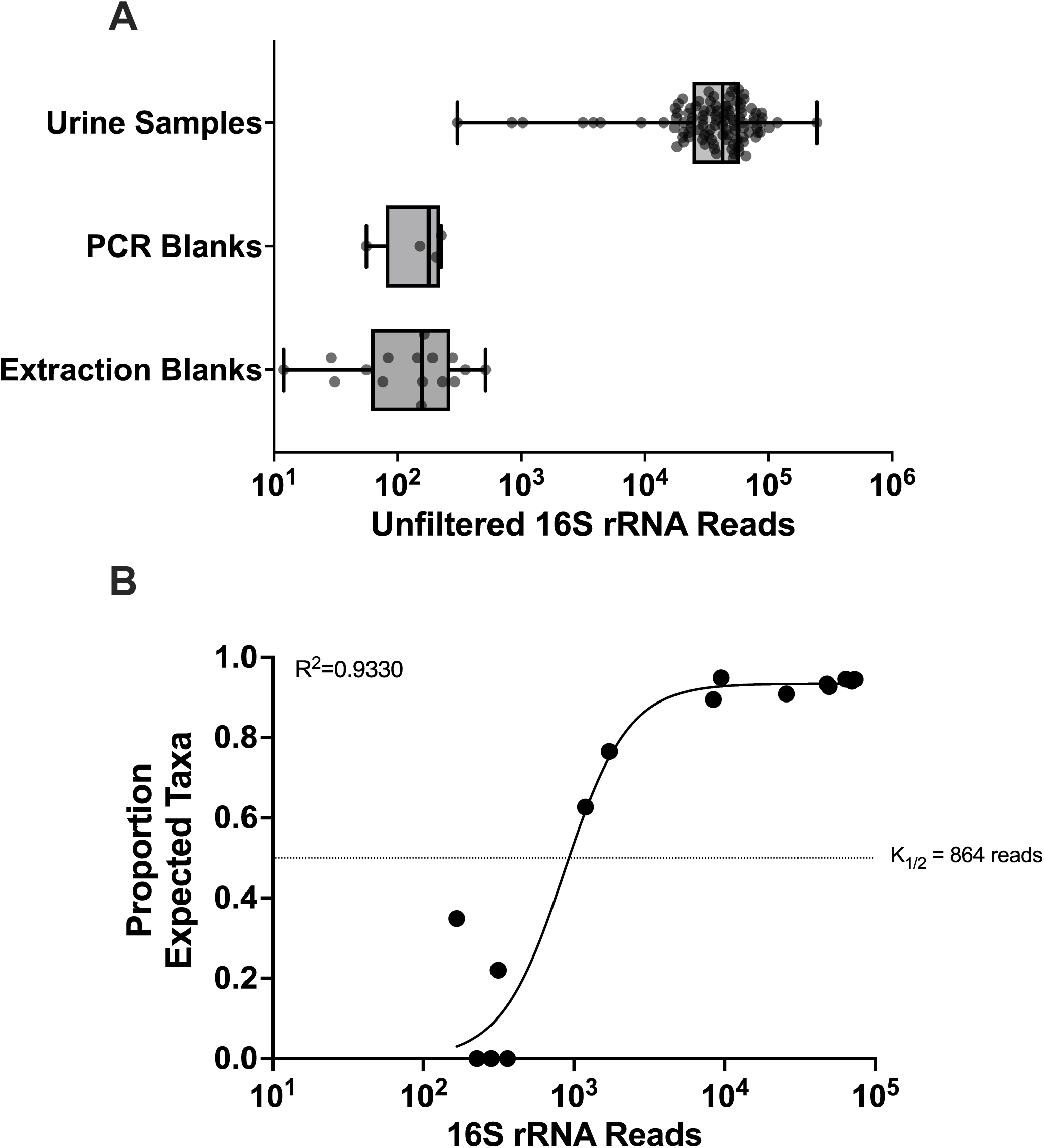
Unfiltered Read Counts and KatharoSeq 16S rRNA Read Threshold. **A)** Read counts of samples and blanks prior to bioinformatic decontamination. The box represents the 25th to 75th percentiles, while the whiskers represent the minimum and maximum read counts. Each data point is a unique sample. **B)** The sequenced taxonomic composition of serial dilutions of the ZymoBIOMICS™ Microbial Community Standard was compared to the known taxonomic composition of the mock community standard. Allosteric sigmoidal non-linear regression was conducted in GraphPad Prism v. 10.2.2.

**Figure S2.**
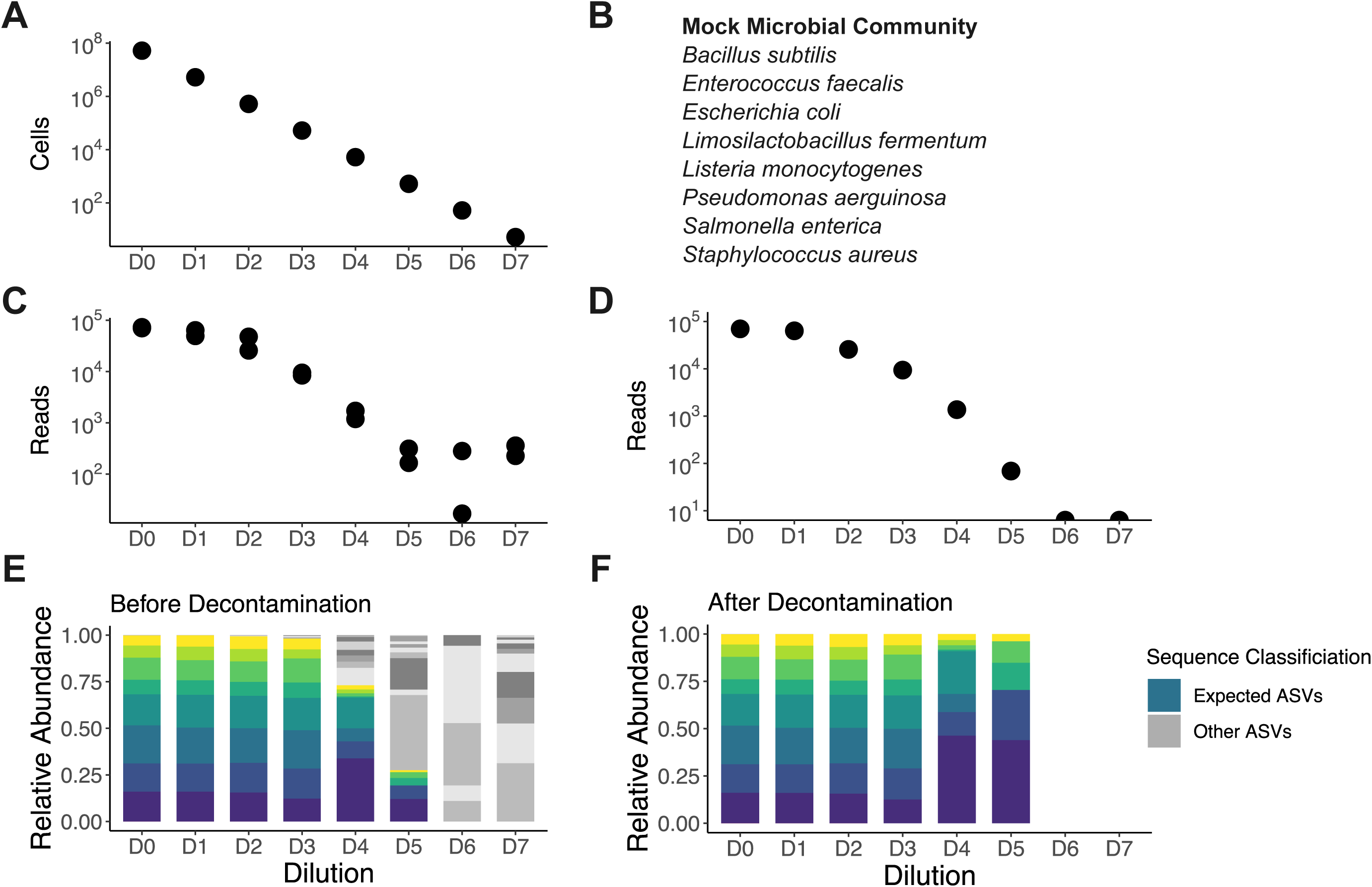
Bioinformatic Decontamination of 16S rRNA Sequencing from a Mock Microbial Community. **A)** Cell count of a ten-fold serial dilution of a mock microbial community. **B)** Bacterial composition of the Zymo D6300 Mock Microbial Community. **C)** Read count of the mock microbial dilution prior to bioinformatic decontamination. **D)** Read count of the mock microbial community after bioinformatic decontamination. **E)** Stacked bar plot of sample composition before decontamination. **F)** Stacked bar plot of sample composition after decontamination. For panels **E - F**, the colored bars match expected taxa from the mock microbial community while the gray bars represent contaminants.

**Figure S3.**
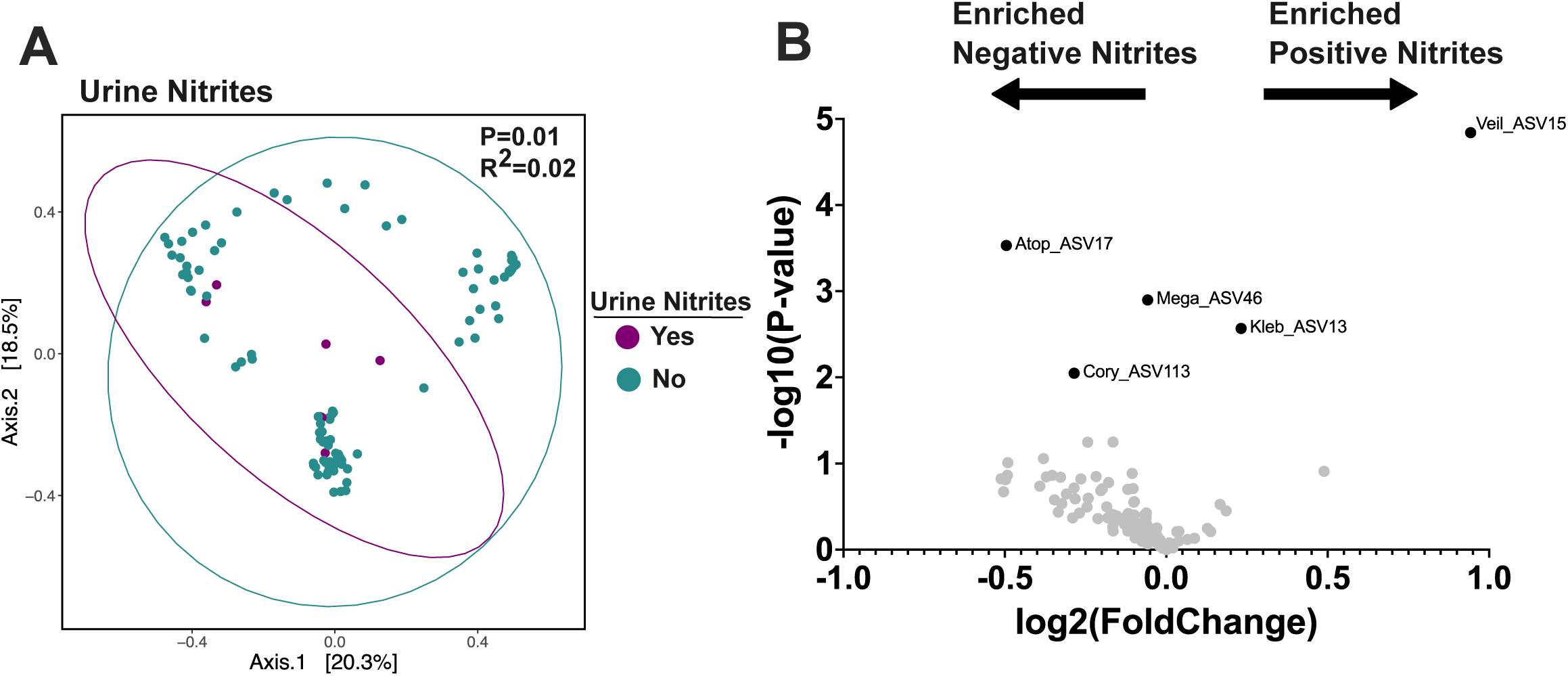
Association of Urinalysis Results with Urobiome Composition. **A)** Principal component analysis of Bray-Curtis distances of samples stratified by presence of urine nitrites. P value calculated by PERMANOVA. **B)** Volcano plot of ASVs with respect to the presence of urine nitrites. Differential abundance analysis conducted with MaAsLin2. Black labeled dots were significantly differentially abundant (MaAsLin2 FDR<0.25).

**Additional file 1. Deidentified Subject Metadata and Unfiltered Read Counts.**

**Additional file 2. ASV Taxonomic Assignment.**

**Additional file 3. MaAsLin2 Differential Abundance with Respect to Menopausal Status.**

**Additional file 4. MaAsLin2 Differential Abundance with Respect to Recent UTI.**

**Additional file 5. MaAsLin2 Differential Abundance with Respect to Pain Localization Phenotype.**

**Additional file 6. MaAsLin2 Differential Abundance with Respect to Overall GUPI.**

**Additional file 7. MaAsLin2 Differential Abundance with Respect to Urinary GUPI.**

**Additional file 8. ASVs Included in Persistence Analysis.**

## DECLARATIONS

### Ethics approval and consent to participate

Study procedures and sample collection were approved by the Vanderbilt University Medical Center IRB (IRB #193461). Subjects provided written informed consent.

### Consent for publication

No individual-level data or identifying details are included in this manuscript.

### Availability of data and materials

Raw sequencing data is publicly available under NCBI BioProject PRJNA1141486. Bioinformatic code is available: https://github.com/reaset41/IC-Urobiome-Longitudinal.

### Authors’ contributions

SAR, MH, and LCM conceptualized the study. LCM, AGK, AMR, and AA collected patient samples and patient clinical data. SAR, MG, JDF, and BTF analyzed the sequencing data. SAR wrote the manuscript with input from all authors.

## Acknowledgements

We thank the patients who participated in this study. We thank the Vanderbilt Clinical Research Center for collecting samples and acquiring patient information. Graphics were created in Inkscape using open-source vector templates from SciDraw.io, bioart.niaid.nih.gov, bioicons.com.

